# scPACE: A Personalized Autoencoder for Cell Evolution in Single-Cell Transcriptomics

**DOI:** 10.64898/2026.11.19.767173

**Authors:** Jingxuan Bao, Hsin-Yu Lai, Michael S. Cuoco, Cindy T.J. van Velthoven, Kyle J. Travaglini, Mariano Ignacio Gabitto

**Affiliations:** Allen Institute Seattle, WA 98109

## Abstract

Single-cell studies increasingly profile large cohorts of individuals, measuring thousands of cells in each donor. These datasets contain information at two levels: variation among individual cells and among donors. Analyzing both levels requires methods that can learn cellular representations, infer cell and donor states, and account for cell types that may be missing in some donors. Existing methods partially address these tasks. We introduce scPACE, a hierarchical personalized variational autoencoder handling these tasks. scPACE learns a cell representation, a cell state, and then aggregates cells within each type of cell to infer a donor state. It also imputes missing cell types with confidence estimates. Across simulated cohorts and a developing mouse visual-cortex dataset, scPACE recovered held-out cell and donor states accurately while achieving efficient cell-type imputation and representation. scPACE provides a unified framework for learning phenotypes across cellular and donor scales in cohort-scale single-cell transcriptomics.

## 1 Introduction

Cohort-scale single-cell transcriptomics links molecular variation among individual cells to phenotypes measured in the donors from whom they are collected. These datasets can contain phenotypes at two levels. Each donor has a phenotype, such as age, disease stage, or treatment, while individual cells can have their own maturity, perturbation, or activity scores. Analyzing such cohorts therefore requires solving several closely connected tasks that are often handled separately.

For data exploration and biological discovery, cell- and donor-level representations must account for relevant technical covariates. At the same time, phenotypes at both levels must be modeled without treating one as a substitute for the other. Assigning the donor phenotype to every cell creates pseudoreplication and cannot identify cells that depart from their donor’s average, while ignoring cell-level measurements discards within-donor variation. These representations must also support label transfer from well-annotated reference data to unlabeled cells and donors. Importantly, these tasks are statistically coupled. The learned representation shapes the cell-type posterior, cell type determines how expression is decoded and how cells contribute to donor summaries, and cell states are nested within donor states. Handling these tasks separately can compound errors and prevents each task from informing the others. A single model that learns them jointly is therefore needed.

Existing methods address individual parts of this problem. For cell-level phenotype association, co-varying neighborhood analysis (CNA) relates sample attributes to neighborhood-abundance variation and maps the resulting effects back to individual cells [Reshef et al., 2022], while MELD smooths condition indicators over a cell graph to estimate relative condition likelihoods [Burkhardt et al., 2021]. Trajectory methods provide related scalar descriptions of cellular progression. Sling-shot fits principal curves through a cluster-level minimum spanning tree without phenotype supervision [Street et al., 2018]. PhenoPath allows observed covariates to modify a shared latent pseudo-time [Campbell and Yau, 2018], while Psupertime and Sceptic use ordered capture labels to learn supervised cell orderings [Macnair et al., 2022, Li et al., 2025]. Deep generative models such as scVI learn batch-aware probabilistic cell embeddings from expression counts [Lopez et al., 2018]. scANVI adds semi-supervised cell-type imputation [Xu et al., 2021], while scVIP uses cell-type-aware multiple-instance learning to connect expression and composition with donor phenotypes [Lai et al., 2026]. However, none of these generative models learns from phenotypes measured at the cell level. Conversely, methods that produce cell-level scores generally use ordered sample labels or condition indicators rather than measured per-cell phenotypes, and lack both a generative count model and a donor latent state. scVIP also requires known cell types, while scANVI includes no phenotype supervision at either level. No existing framework therefore spans all these tasks. The remaining gap is a model that jointly learns cell representations, donor and cell phenotypes, and cell-type transfer.

To address this gap, we introduce *scPACE*, a Personalized Autoencoder for Cell Evolution that jointly solves these tasks within a unified framework. A negative-binomial observation model connects latent cell embeddings to raw expression counts. scPACE learns four linked quantities: a vector representation for each cell, a scalar donor state supervised by donor-level phenotypes, a scalar cell-evolution state nested within the donor state and supervised by cell-level phenotypes, and a cell-type distribution for annotating unlabeled cells. The principal contribution of scPACE is a joint modeling strategy that integrates representation learning, donor-level supervision, cell-level supervision, and cell-type transfer with quantified confidence. We evaluate scPACE across a 12-cohort simulation and 865,012 developing mouse visual-cortex profiles. scPACE achieved the best recovery of donor- and cell-level phenotypes on query samples, outperformed scANVI in cell-class imputation, and matched major representation-learning models in donor mixing.

## 2 Methods

Let cell *n* carry raw counts 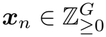 over *G* genes, a batch label *b*_*n*_, a donor index *d*(*n*), and a cell type *c*_*n*_ ∈ {1, …, *K*} that may be unobserved. Let a donor carry a phenotype *y*_*d*_ and optionally, a cell phenotype *y*_*n*_ which can be missing independently. For each cell, scPACE returns one output per task: a vector representation ***z***_*n*_, a scalar cell state *s*_*n*_, a scalar donor state *t*_*d*_ shared by all cells of donor *d*, and a cell-type posterior *q*_*ϕ*_(*c*_*n*_ | ***z***_*n*_), the probability assigned to each of the *K* candidate types given that cell’s representation, where *ϕ* collects the parameters of the amortized inference networks introduced below. Phenotypes serve only as supervision targets and are never read by the encoder. The cell representation ***z***_*n*_ and cell state *s*_*n*_ are inferred from counts, whereas the donor state *t*_*d*_ additionally uses donor label and observed or confidently transferred cell types. Batch enters only the count decoder, while *y*_*d*_ and *y*_*n*_ enter exclusively through supervised losses and are not encoder inputs. Inference therefore needs no phenotype, so a fitted model can score donors and cells whose labels are missing, held out, or not yet measured. Figure 1 summarizes cell-level inference, donor aggregation, the type-aware count model, and the reported outputs. An implementation of scPACE is available at https://github.com/bao96-bioinfo/scPACE/tree/NeurIPS26.

**Figure 1:**
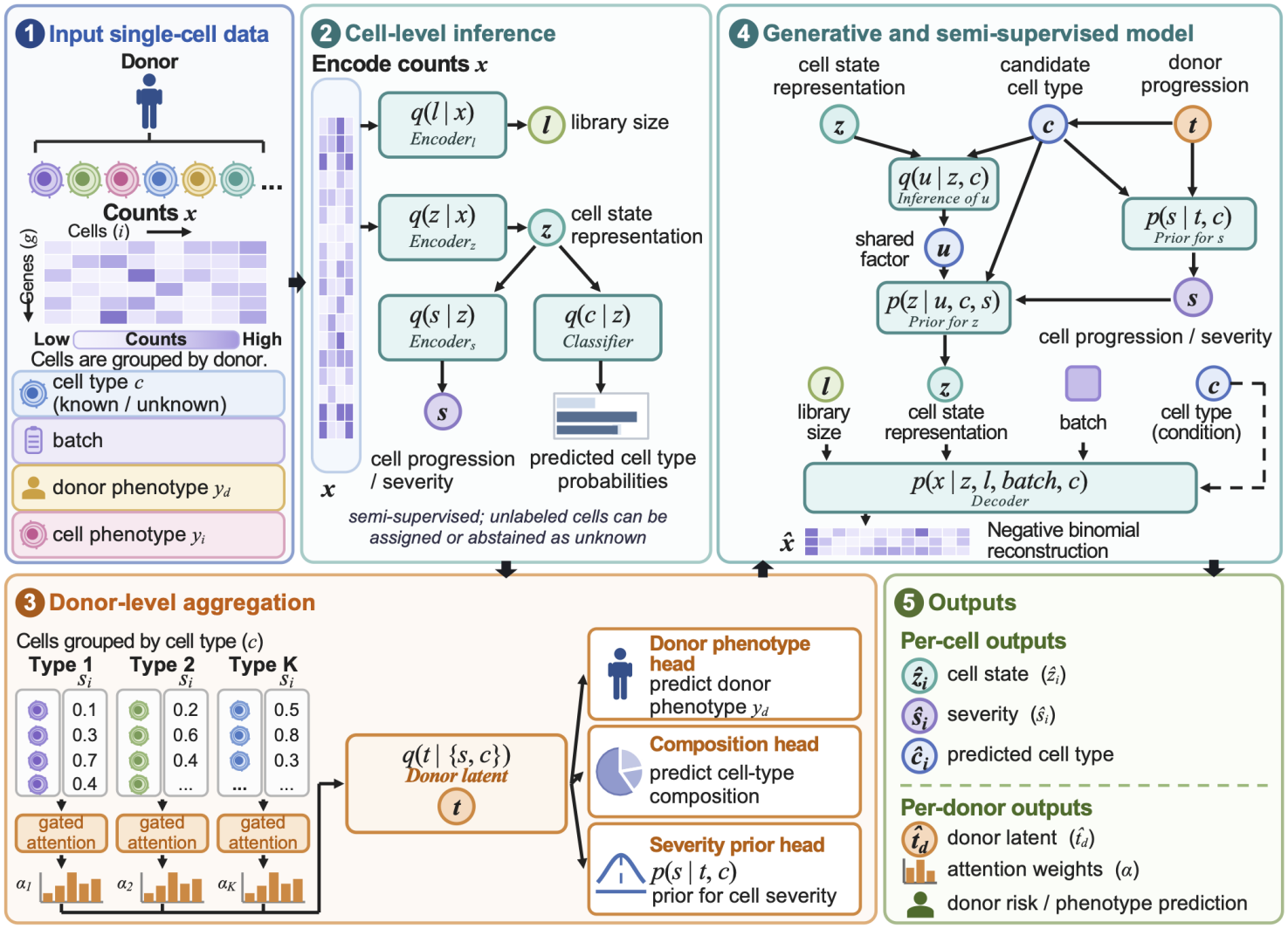
Overview of scPACE. scPACE jointly learns cellular representations, cell states, cell types, and donor phenotypes. (1) Counts are organized by donor with batch, cell-type, and phenotype information. (2) Cell encoders learn representations, scalar cell states, and cell-type probabilities. (3) Type-specific MIL attention aggregates cells to infer donor states and phenotypes. (4) A cell-type- and batch-aware decoder reconstructs expression. (5) scPACE outputs cell and donor states, cell-type predictions with confidence, attention summaries, and phenotype predictions.

scPACE uses a hierarchical generative model that preserves the donor-cell structure of cohort data. Each donor *d* is assigned a latent state *t*_*d*_. Conditional on this donor state, the model draws the cell type *c*_*n*_ and scalar state *s*_*n*_ of each cell *n* from that donor. Therefore, cells from the same donor can differ from one another while still reflecting a common donor-specific context. At the cell level, an auxiliary variable ***u***_*n*_, the cell type *c*_*n*_, and the scalar state *s*_*n*_ jointly determine the distribution of the cell representation ***z***_*n*_. The decoder then combines this representation with the onehot embedding of cell-type onehot(*c*_*n*_) and batch onehot(*b*_*n*_) to obtain the relative expression levels ***ρ***_*n*_. Finally, the library size *ℓ*_*n*_ scales these relative expression levels to produce the expected counts. Thus, ***z***_*n*_ and *c*_*n*_ capture biological variation, while *b*_*n*_ and *ℓ*_*n*_ account for batch and sequencing-depth effects:

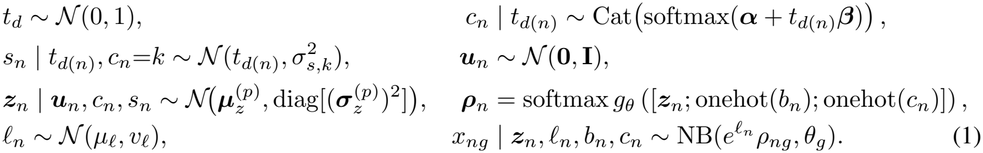

Here, *d*(*n*) identifies the donor of cell *n*. The parameters ***α*** and ***β*** determine the baseline cell-type composition and its dependence on the donor state, while 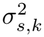 describe the residual variation of cell state within type *k*. The neural prior maps ***u***, *c*, and *s* to 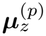 and 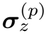, the mean and standard deviation of the conditional distribution of ***z***_*n*_. In the count model, 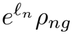 is the expected count for gene *g* in cell *n, θ*_*g*_ is its gene-specific inverse dispersion, and (*µ*_*ℓ*_, *v*_*ℓ*_) are estimated from the log library sizes in the training data. Because *t*_*d*_ controls both *p*(*c*_*n*_ | *t*_*d*_) and the center of the cell-state distribution, scPACE can represent donor-associated changes in cell-type composition together with expression changes within each cell type. Batch and cell type enter the decoder as separate one-hot covariates, allowing their contributions to expression to be modeled separately.

The inference network factorizes through

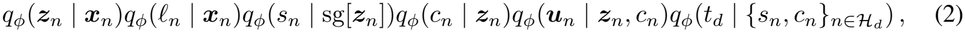

where *ℋ* _*d*_ contains cells with observed cell types, and sg denotes the stop-gradient operator. A donor is a variable-size bag. To infer the donor state, scPACE first summarizes cells separately within each cell type. A gated-attention mechanism assigns larger weights to cells that are more informative and computes a weighted summary for each type [Ilse et al., 2018]. The model combines these cell-type-specific summaries with the proportion of each cell type in the donor. Two neural networks then estimate the mean and variance of *q*_*ϕ*_(*t*_*d*_), representing the donor state and its uncertainty.

Because the summaries are weighted sums, the result does not depend on the order of the cells. For a cell with unknown type, scPACE enumerates all *K* categories and averages the nuisance posterior, representation prior, and reconstruction with weights *q*_*ϕ*_(*c*_*n*_ = *k* | *z*_*n*_). The cell therefore contributes to count modeling without a fabricated label. During training, donor pooling and type-dependent losses use observed cell types only. During testing, donor pooling also uses transferred types whose confidence exceeds the threshold *T* . Supplementary Section A.1 gives the complete factorization and exact marginalization.

To orient both scalar states toward increasing phenotype, each supervision head uses the positive-slope parameterization *w*^+^ = softplus(*w*) *>* 0:

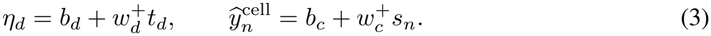

The donor loss is binary cross-entropy on logit *η*_*d*_ for a binary outcome or squared error for a continuous outcome. The cell loss is 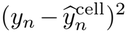 over every finite cell target, including cells whose types are unknown. Training minimizes the weighted objective,

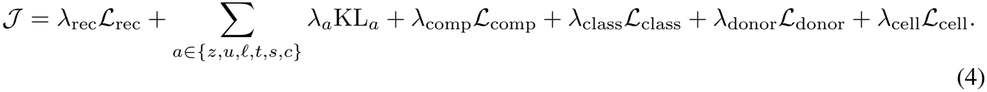

## 3 Simulation experiments

We generate 12 matched cohorts by crossing two expression regimes with *D*∈ { 5, 30, 100} donors and *N*_*d*_ ∈ {100, 1000} cells per donor. In Regime A, cell state 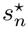 affects expression through smooth linear and quadratic gene programs that are partly shared and partly cell-type specific. Regime B retains the same donors, structural truths, phenotypes, cell types, nuisance variables, and data partitions, but reduces the smooth program and adds an 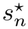 -dependent, cell-specific rough component. Consequently, transcriptomically neighboring cells are less likely to have similar states in Regime B, making it more challenging without changing the underlying truths or supervision. Donor truth 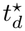 controls a noisy continuous donor phenotype and type composition, while nested cell truth 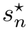 controls a noisy cell phenotype and type-specific count programs. We hold out 20% of donors before fitting and compare scPACE with six scalar-state methods (Sceptic, psupertime, PhenoPath, Sling-shot, CNA, and MELD) and three representation models (scVI, scANVI, and scVIP). All methods use identical training donors, while ground truths and query phenotypes remain evaluation-only. Supplementary Section A.6 provides the complete generator, partitions, training profile, and metrics.

Across the query donor benchmark, scPACE is the only evaluated method providing all four output classes: a native donor state, a native cell score, cell-type imputation, and a reusable representation. It achieves donor Pearson *r* = 0.838 over the eight cohorts with estimable donor correlations. Across all 12 cohorts, it achieves calibrated donor RMSE = 0.368, cell Pearson *r* = 0.903, calibrated nRMSE = 0.435, and cell-type imputation accuracy = 0.981. The four learned representations have similar within-type donor-mixing ratios (0.982-0.985), close to the random-mixing reference of 1. Table 1 reports all methods and cohort-level variability.

**Table 1:** Performance on query donors across 12 simulated cohorts. Each entry reports the equally weighted mean (standard deviations) across cohorts. Donor Pearson correlation is averaged eight cohorts because four simulated cohorts contain only one query donor.

|  | Native donor output vs. $t^\star$ |  | Reported cell score vs. $s^\star$ |  | Native type | Donor mixing |
|---|---|---|---|---|---|---|
| Method | Pearson *r* ↑ | RMSE↓ | Pearson *r* ↑ | nRMSE↓ | Accuracy↑ | kNN/random↑ |
| **scPACE** | **0.838 (0.302)** | **0.368 (0.165)** | **0.903 (0.010)** | **0.435 (0.024)** | **0.981 (0.009)** | 0.982 (0.008) |
| scVIP | 0.767 (0.314) | 0.632 (0.273) | — | — | — | 0.985 (0.008) |
| scANVI | — | — | — | — | 0.928 (0.151) | 0.985 (0.008) |
| scVI | — | — | — | — | — | 0.985 (0.008) |
| Sceptic | — | — | 0.222 (0.199) | 0.977 (0.058) | — | — |
| psupertime | — | — | 0.162 (0.327) | 0.979 (0.091) | — | — |
| PhenoPath | — | — | -0.520 (0.056) | 0.893 (0.059) | — | — |
| Slingshot | — | — | -0.016 (0.388) | 0.947 (0.094) | — | — |
| CNA | — | — | 0.218 (0.021) | 0.979 (0.005) | — | — |
| MELD | — | — | 0.228 (0.148) | 0.990 (0.055) | — | — |

## 4 Real-data experiments

We evaluated scPACE on 865,012 mouse visual-cortex profiles where 568,654 class-annotated cells are from the Gao et al. developmental atlas and 296,358 project-processed nuclei are from GSE190940, which samples postnatal ages and visual-rearing conditions [Gao et al., 2025, Cheng et al., 2022]. Chronological age was normalized without external-query units, and was the donor target. Because no cell-level maturity measurement was available, we used a prespecified mature-versus-progenitor AUCell-like signature as an expression-derived proxy for cellular maturation. Unlike donor age, which is shared by all cells within a specimen, this rank-based score captures within-specimen variation and provides a continuous, interpretable target across cohorts. Accordingly, *s* is interpreted as relative signature-aligned maturity. GSE190940 class labels were treated as missing. We reserved 21 complete donor/sample units (183,656 cells) for external evaluation, then divided the reference units into 61 training and 6 internal-validation units. All methods used the same training units, although the baselines followed their native donor-level or unsupervised training contracts and did not receive scPACE’s direct cell-maturity loss. Supplementary Section A.7 gives the complete construction and metric definitions. On external-query cells, scPACE *ŝ* correlated with the maturity proxy at *r* = 0.919; the six benchmark scores ranged from −0.060 to 0.438 (Fig. 2a). The native scPACE donor state tracked observed normalized age across the 21 query units (*r* = 0.942), whereas the separately defined scVIP donor-phenotype prediction reached *r* = 0.542 (Fig. 2b). Raw class-imputation accuracy on 52,814 query Gao cells was 0.977 for scPACE and 0.976 for scANVI (Fig. 2c), indicating comparable overall accuracy. Within-class donor-mixing ratios were also similar across the four representations (0.132-0.139) (Fig. 2d). The shared scPACE latent UMAP organized broad cell classes and displayed a continuous *ŝ* gradient (Fig. 2e,f). Because GSE190940 lacks class ground truth, its class colors are transferred predictions.

**Figure 2:**
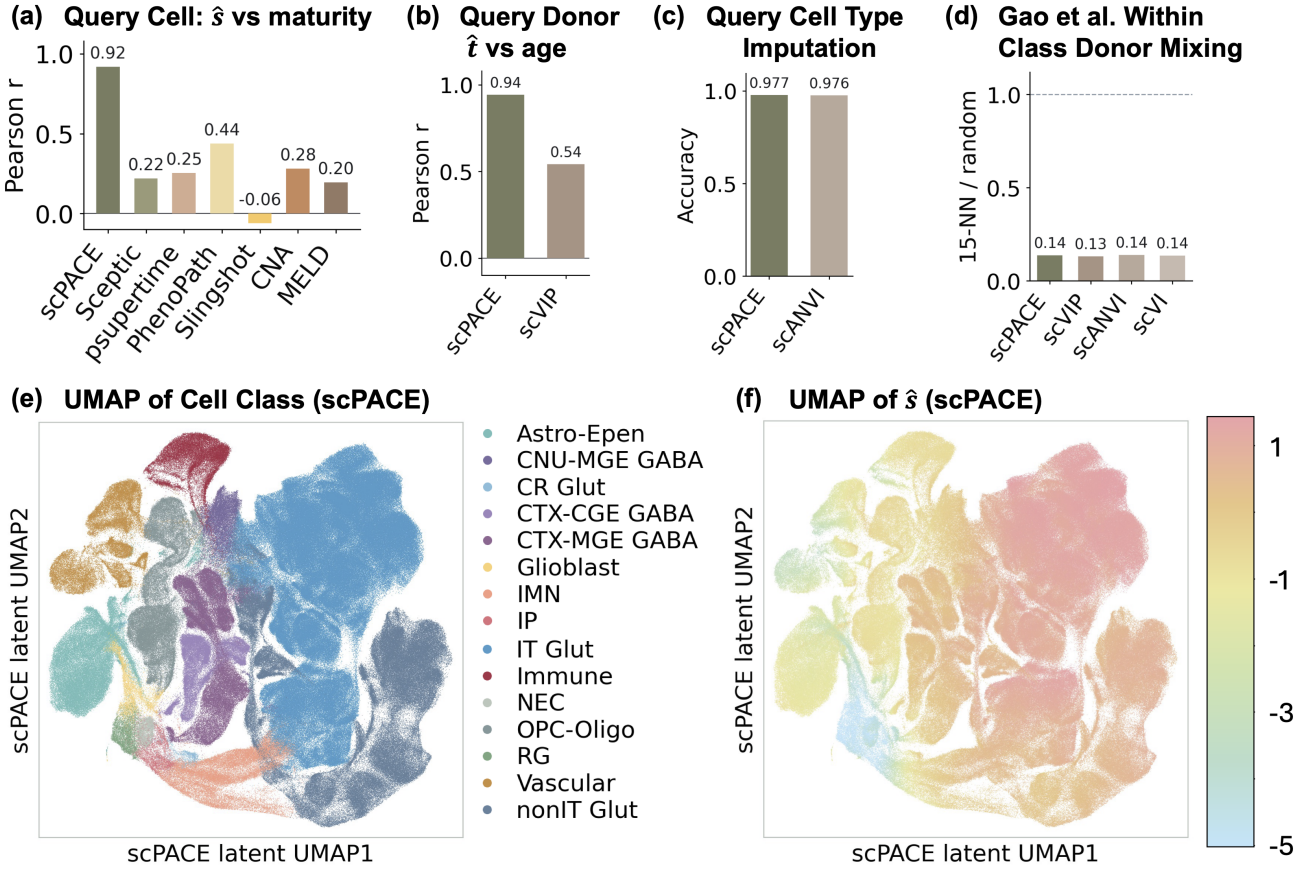
Evaluation on query visual-cortex data. (a) Pearson correlation between each method’s cell-level output and the predefined expression-derived maturity target across 183,656 query cells. (b) Pearson correlation with observed normalized age across 21 query units. (c) Raw argmax cell-class accuracy across 52,814 typed Gao query cells. (d) Within-class 15-nearest-neighbor mixing across donors. A value of 1 indicates random mixing. (e,f) scPACE latent space UMAP, colored by cell class in (e) and *s* in (f). Cell-class labels are observed for Gao et al. and predicted for GSE190940. Bars show single-experiment estimates.

## 5 Conclusion and limitations

scPACE provides a unified model for cell representation, scalar cell and donor states, type-aware donor pooling, and label transfer. Across simulations on query donors, it recovered both cell- and donor-level structure while retaining annotation and representation performance. In the real-data study, scPACE achieved stronger held-out cell-state and donor-phenotype recovery than the evaluated baselines while matching scANVI’s overall cell-type-imputation accuracy. Although our AUCell-like maturation target provides an interpretable weak-supervision signal when no measured cell phenotype is available, it is not an independent biological endpoint and should be viewed as an example of cell-level supervision. It is constructed from a small prespecified marker gene set in the same expression matrix provided to the model and may therefore reflect marker selection. Its recovery demonstrates transfer of a signature-aligned relative state rather than independent validation of absolute cellular maturity. Future work should incorporate independent or disease-relevant cellular phenotypes.

## A Supplementary Materials

### A.1 Full probabilistic and variational factorization

For donor *d*, conditional on observed batches, the joint distribution represented by Eq. (1) is

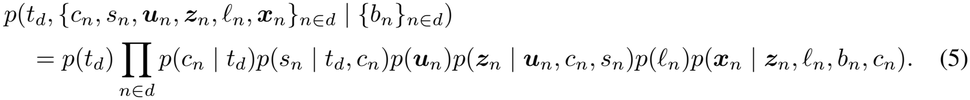

Let *ℋ*_*d*_ and *U*_*d*_ index hard-typed and unknown-type cells. The semi-supervised approximation is

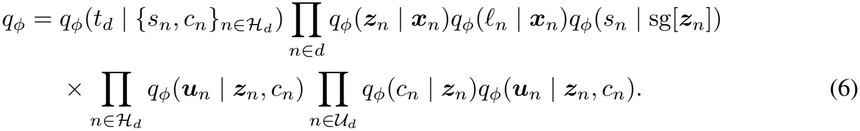

All continuous factors are diagonal Gaussians parameterized by neural networks, and *q*_*ϕ*_(*c*_*n*_| ***z***_*n*_) is categorical. Neither *y*_*d*_ nor *y*_*n*_ appears in a conditioning set. Labels act only through the supervised losses.

For unknown *c*_*n*_, let *F* (*c*_*n*_) denote any per-cell term of the objective whose value depends on the type. Three such terms occur, including the nuisance KL, the conditional representation KL, and the negative-binomial reconstruction,

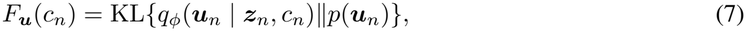

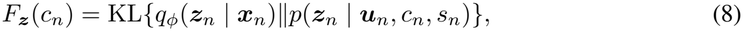

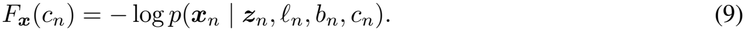

Each is evaluated exactly as

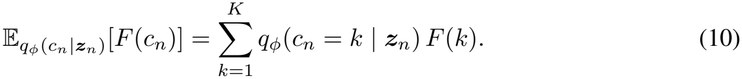

### A.2 Objective, missing labels, and gradient routing

We define a bag as the set of cells of a donor *d*. The sets *ℋ*_*d*_ and *U*_*d*_ index its hard-typed and unknown-type cells, and *N*_*d*_ = |*H*_*d*_| + |*U*_*d*_| is the full bag size.

#### Reconstruction loss

Under the mean-inverse-dispersion parameterization

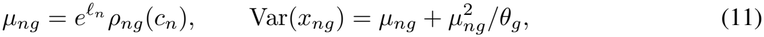

the per-cell reconstruction term is the gene-summed negative log-likelihood of the count model, 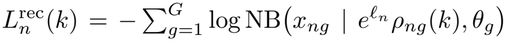. Its contribution to *ℒ*_rec_ is 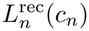 for a hard-typed cell and the type expectation 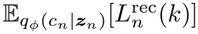 of Eq. (10) for an unknown cell, averaged over the bag.

#### KL loss

The five Gaussian divergences KL_*z*_, KL_*u*_, KL_*ℓ*_, KL_*t*_, KL_*s*_ pair diagonal-Gaussian variational and prior factors and are therefore evaluated in closed form. Two of these terms depend on the assumed cell type: KL_*u*_ through its posterior *q*_*ϕ*_(***u***_*n*_| ***z***_*n*_, *c*_*n*_) and KL_*z*_ through its prior *p*(***z***_*n*_ |***u***_*n*_, *c*_*n*_, *s*_*n*_). On a hard-typed cell, each is evaluated at the observed type; and on an unknown cell, each is averaged over the *K* candidate types by the type expectation of Eq. (10), using the same classifier weights *q*_*ϕ*_(*c*_*n*_=*k*| ***z***_*n*_) that weight the reconstruction term. The categorical term KL_*c*_ applies only to unknown cells and regularizes the classifier *q*_*ϕ*_(*c*_*n*_| ***z***_*n*_) toward the donor composition. When a donor has hard-typed cells (*ℋ*_*d*_ ≠ ∅), the composition model is estimable and serves as the detached target,

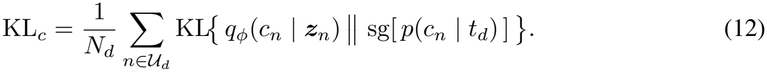

When a donor has no hard-typed cells (*ℋ*_*d*_ = ∅), the composition is unanchored and the target falls back to the uniform prior,

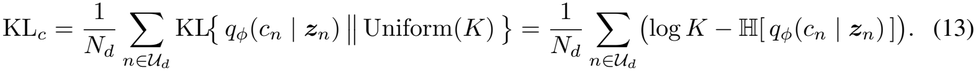

#### Cell-type supervision loss

Two terms supervise cell type on hard-typed cells. The classification loss is the cross-entropy of the classifier,

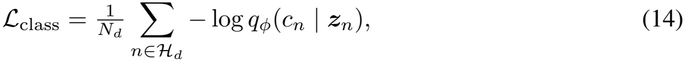

and the composition loss is the negative log-likelihood of the donor-composition model,

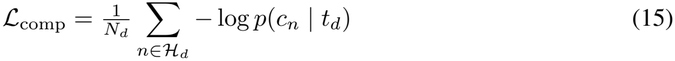

With

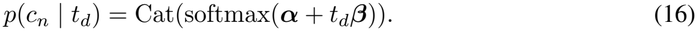

Both sums run over *ℋ*_*d*_ but are normalized by the full bag size *N*_*d*_.

#### Phenotype supervision and missing labels

Donor and cell phenotypes supervise the positive-slope heads of Eq. (3). ℒ_donor_ is a binary cross-entropy on the donor logit *η*_*d*_ for a binary outcome or a squared error for a continuous outcome, and ℒ_cell_ is the mean squared error 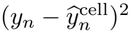 over cells with a finite target. The donor phenotype loss and the donor KL_*t*_ are donor-level scalars. Every supervised term masks nonfinite targets.

#### Gradient routing

The generative model in Eq. (1) specifies how the latent variables depend on one another: the donor state *t*_*d*_ shapes each cell’s type *c*_*n*_ and state *s*_*n*_; these, with the auxiliary variable ***u***_*n*_, shape the representation ***z***_*n*_; and ***z***_*n*_, with library size and batch, generates the counts. Training does not send a gradient back through every one of these dependencies. Two stop-gradient operators sg[· ] restrict gradient flow through a subset of those parameters. First, the cell-state encoder reads a detached representation, *q*_*ϕ*_(*s*_*n*_ | sg[***z***_*n*_]). Therefore, neither the cell-state supervision *ℒ*_cell_ nor KL_*s*_ propagates into the representation encoder along the cell-state route. Second, the composition prior is detached inside KL_*c*_ (the target *π*_*d*_ above), so KL_*c*_ updates the classifier *q*_*ϕ*_(*c*_*n*_ | ***z***_*n*_) rather than the composition model *p*(*c*_*n*_ | *t*_*d*_), which is trained solely by ℒ_comp_.

### A.3 Identifiability and interpretation

Some latent quantities in scPACE are identifiable only up to transformations, and supervision is what removes the residual freedom. The representation ***z***_*n*_ is not identifiable coordinate by coordinate, which means a rotation, or any other transformation the decoder can absorb, gives the same likelihood. Therefore, no individual axis of ***z***_*n*_ carries intrinsic meaning. The donor state *t*_*d*_’s standard-Normal prior fixes a reference location and scale. When donor supervision is active, the positive-slope donor head fixes its sign so that larger *t*_*d*_ corresponds to higher donor phenotype. The cell state *s*_*n*_ is oriented the same way by the positive-slope cell head, but only when finite cell labels and a nonzero cell-loss weight are present.

### A.4 Baseline models and formulations

For each baseline below, we describe the method as originally published and then note separately any changes we made so it could score query donors.

#### A.4.1 Sceptic

Sceptic is a supervised pseudotime method for data in which cells carry an ordered capture label *Y*_*n*_ ∈ {1, …, *R }*, where *R* is the number of stages and larger values denote later stages [Li et al., 2025]. For each cell, it returns a scalar pseudotime that places the cell along this ordered axis.

In their published method, each cell *n* is represented by a feature vector ***r***_*n*_. For every stage *r*, Sceptic trains a binary one-versus-rest support-vector machine (SVM) that separates the cells of stage *r* from all other cells using a radial-basis-function (RBF) kernel

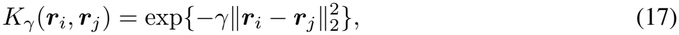

in which ∥·∥_2_ is the Euclidean norm and *γ >* 0 is the kernel bandwidth. The decision score of the stage-*r* machine at a point ***r*** is

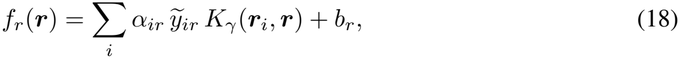

where the sum runs over training cells *i*; 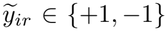 marks whether training cell *i* belongs to stage *r*; *α*_*ir*_ ≥ 0 are the learned dual coefficients, which is nonzero only on the support vectors; and *b*_*r*_ *b* is the bias. The *R* decision scores are calibrated to class probabilities *p*_*nr*_ = Pr(*Y* _*n*_ = *r* | ***r*** _*n*_ ), and the reported pseudotime is their probability-weighted mean

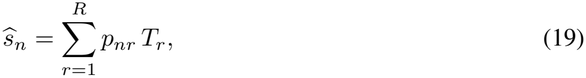

where *T*_*r*_ is the numeric value assigned to stage *r*, i.e. its representative position on the phenotype axis.

For computational scalability and compatibility with the held-out-donor workflow, we use Sceptic’s published GPU-enabled XGBoost option rather than its default RBF-kernel SVM. We refer to this implementation as *adapted Sceptic*. Specifically, each cell is represented by principal-component (PCA) coordinates ***r***_*n*_, with the PCA basis learned from training donors only and then applied to all cells. In place of the kernel SVM, we use the classifier-construction routine shipped with the Sceptic package, which builds a gradient-boosted-tree (XGBoost) classifier, and we train it on the trainingdonor cells’ PCA coordinates and stage labels. When the phenotype is continuous (e.g. donor age), we form the *R* ordered stages by quantile-binning the phenotype over the unique training donors.

Query-donor phenotypes never influence the model training, and the original continuous phenotype is kept for evaluation. Each *T*_*r*_ is the median phenotype among the training-donor cells assigned to stage *r*. Finally, the trained classifier predicts stage probabilities *p*_*nr*_ for all cells, including those query donors, and the reported pseudotime is 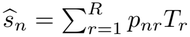.

#### A.4.2 psupertime

psupertime learns a sparse linear ordering of cells from sequential cell labels [Macnair et al., 2022]. In their published method, each cell *n* is represented by a standardized log-expression vector ***g***_*n*_ over the highly variable genes and carries an ordered label *Y*_*n*_ ∈{1, …, *R* }, where *R* is the number of stages. psupertime fits a proportional-odds model, which gives the probability that a cell falls at or below stage *r* as

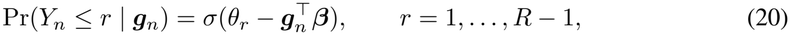

where *σ*(*x*) = 1*/*(1 + *e*^*−x*^) is the logistic function, *θ*_1_ *<*· · · *< θ*_*R* −1_ are ordered cut points that separate consecutive stages, and ***β*** is a gene-weight vector shared across all stages. It estimates ***β*** and the cut points by minimizing the joint ordinal negative log likelihood together with an *L*_1_ penalty *λ*∥***β***∥_1_, whose strength *λ* controls sparsity. The resulting pseudotime is the linear score 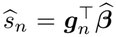, and the genes with nonzero 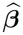 are those whose expression changes coherently with the label sequence.

Because psupertime already learns a linear coefficient vector, its score extends to new cells directly, so our only changes concern how it is trained for donor-held-out evaluation. When the phenotype is continuous (e.g. donor age), we form the *R* ordered stages by quantile-binning the phenotype over the unique training donors, so each training donor contributes equally to the bin edges, and we keep the original continuous phenotype for evaluation. We then fit the model to the training-donor cells’ standardized log-expression with regularization fixed at *λ* = 0.01. Finally, the learned coefficient vector 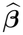 scores every cell, including those of query donors through 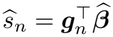.

#### A.4.3. PhenoPath

PhenoPath is a Bayesian latent-factor model that learns a cell trajectory whose shape can vary with observed covariates [Campbell and Yau, 2018]. In their published method, let *g*_*ng*_ be the normalized expression of gene *g* in cell *n*, ***v***_*n*_ a covariate vector for that cell, and *z*_*n*_ its latent pseudotime. The generative model is

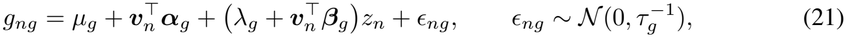

where *µ*_*g*_ is the baseline expression of gene *g*, ***α***_*g*_ are its covariate loadings, *λ*_*g*_ is its shared-trajectory loading, ***β***_*g*_ are its covariate-trajectory interaction loadings, and *ϵ*_*ng*_ is Gaussian noise with gene-specific precision *τ*_*g*_. The three structured terms play distinct roles: 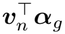 is expression that depends on the covariate alone, *λ*_*g*_*z*_*n*_ is a trajectory shared across cells, and 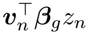 is the part of the trajectory that the covariate modulates. Gaussian-Gamma priors shrink these parameters, with sparse shrinkage on ***β***_*g*_. PhenoPath applies coordinate-ascent mean-field variational inference to estimate the posterior trajectory and interaction effects.

We fit PhenoPath to the normalized variable-gene expression of the training cells alone, using the training-donor phenotype as the modeled covariate ***v***_*n*_, and we take each training cell’s score to be the posterior pseudotime *z*_*n*_. PhenoPath returns a pseudotime only for the cells in its training set and cannot directly score a new cell. To score a query cell, we therefore borrow the pseudotimes of nearby training cells. We first place all cells in a PCA space whose axes are learned from the training cells. For each query cell, we then find its *k* nearest training cells in that space and set its score to the average of their PhenoPath pseudotimes, weighting each neighbor by the inverse of its distance so that closer training cells count more. The number of neighbors *k* and this weighting are fixed in advance and use no held-out information.

#### A.4.4 Slingshot

Slingshot is an unsupervised trajectory method [Street et al., 2018]. In their published method, it works in two stages. First, it recovers the branching structure. It starts from the cell coordinates ***r***_*n*_ in the embedding together with a clustering of the cells, connects the cluster centers into a minimum spanning tree. Then, from a designated starting cluster, it reads off each path through the tree to an endpoint cluster as one candidate lineage. Therefore, a lineage is an ordered chain of clusters that a cell may follow. Second, it lays a smooth curve along each lineage. For lineage *l*, it fits a principal curve ***c***_*l*_(*τ* ), a smooth curve through the middle of that lineage’s cells parameterized by arc length *τ* . Slingshot fits the curve by repeatedly projecting each cell onto the current curve and then re-smoothing the curve through those projections. To place cell *n* on lineage *l*, Slingshot finds the point of the curve nearest the cell and takes the cell’s pseudotime 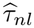 to be that point’s arc length, that is, how far along the curve it lies,

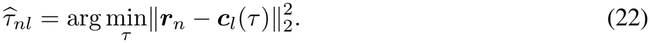

Therefore, a cell near the start of the curve receives a small pseudotime and a cell near the end receives a large one. The lineages are fitted jointly, so their curves coincide along the shared early portion of the trajectory and separate only after a branch point. Each cell also receives lineage weights that say how strongly it belongs to each curve.

We compute the PCA embedding and the clusters from training cells only, fit the cluster tree and the joint curves on those cells, and keep both fixed. To score a query cell, we use Slingshot’s own prediction operator to project it onto these fixed curves and read off its arc-length position, exactly as for a training cell. A cell can lie on more than one lineage, so when it has a pseudotime on several curves we report the average of those pseudotimes, weighted by the cell’s membership in each lineage. No phenotype is used at any point. Slingshot therefore measures geometric position along an inferred lineage and does not produce a donor-phenotype predictor or a probabilistic count representation.

#### A.4.5 CNA

Co-varying neighborhood analysis (CNA) tests whether donors with different phenotypes have different cell states defined by gene-expression similarity [Reshef et al., 2022]. In their published method, CNA first builds a graph that connects each cell to its nearest neighbors in expression space. A neighborhood is a small local region of this graph, the cloud of transcriptionally similar cells around one anchor cell. For each donor and each neighborhood, CNA measures an abundance, which is how densely that donor’s cells populate that neighborhood. These abundances form the neighborhood-abundance matrix *A* ∈ ℝ^*D×J*^ for *D* donors and *J* neighborhoods. As the phenotype varies across donors, the abundances of many neighborhoods rise and fall together. CNA extracts this shared pattern with a truncated singular-value decomposition of *A*, then tests whether it is associated with the phenotype,

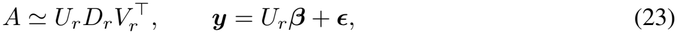

where each retained component represents a pattern of cell-neighborhood abundance across donors. The corresponding column of *U*_*r*_ gives each donor’s score for that pattern, the associated value in *D*_*r*_ measures the strength of the pattern, and the corresponding column of *V*_*r*_ shows how much each cell neighborhood contributes to it. ***y*** is the donor phenotype; ***β*** are the regression coefficients on those patterns; and ***ϵ*** is noise. The regression tests for a global association between the phenotype and these patterns. Mapping 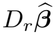 back through *U*_*r*_ gives a coefficient for each neighborhood, which CNA assigns to individual cells according to their neighborhoods. CNA is therefore a donor-level abundance-association method whose output is a localized per-cell score.

We build the cell graph and the neighborhood-abundance matrix from training cells and supply only training-donor phenotypes. CNA’s native coefficients are available for those training cells, and we score query cells by the common inverse-distance PCA *k*-nearest-neighbor projection. Our benchmark implementation of CNA requires at least ten training donors, so it is reported as not estimable for the five-donor simulations, where the prescribed split leaves only three training donors.

#### A.4.6 MELD

MELD estimates how strongly each cell is associated with an experimental condition [Burkhardt et al., 2021]. In their published method, MELD builds a cell-state graph, a graph whose vertices are cells and whose edges join each cell to its nearest neighbors in gene-expression space. For each condition *r*, MELD starts from an indicator over the cells, ***e***_*r*_, which is 1 for cells from condition *r* and 0 otherwise. MELD then smooths it over the graph. Each cell’s value becomes an average of its neighbors’ values, so the estimate changes gradually from cell to cell. This smoothing gives a per-cell value 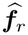 for condition *r*,

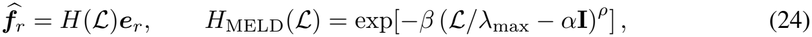

where *ℒ* is the graph Laplacian, and *H*(*ℒ* ) is the smoothing operator. The parameters *α, β*, and *ρ* specify the MELD graph filter, and we retain the software defaults for all three. At each cell, MELD then rescales these smoothed values so that they sum to one across the *R* conditions,

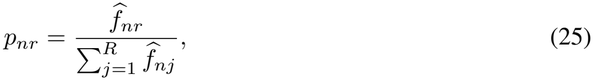

so *p*_*nr*_ is the relative likelihood that cell *n* belongs to condition *r*. For ordered classes, we report *Ŝ* _*n*_ = Σ _*r*_ *p*_*nr*_*T*_*r*_, where *T*_*r*_ is the training-donor median phenotype of class *r*.

We construct the graph, the ordered classes, and the *T*_*r*_ values from training cells and training donors only. Native MELD likelihoods are computed on this graph, and the common training-derived PCA *k*-nearest-neighbor map extends the scalar score to query cells.

#### A.4.7 scVI

scVI is an unsupervised variational autoencoder that learns a low-dimensional representation of each cell from its raw counts while accounting for batch effects [Lopez et al., 2018]. In their published method, each cell *n* has two latent variables: a biological latent ***z***_*n*_∼ *N* (**0**, *I*), which is the low-dimensional representation, and a library-size latent *ℓ*_*n*_, which captures the cell’s overall count depth.

Given these and the cell’s batch *b*_*n*_, its gene counts follow a zero-inflated negative binomial (ZINB) likelihood, a count distribution that also allows extra zeros,

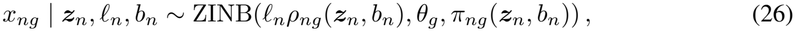

where the rate 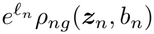 combines the library size with a normalized expression profile ***ρ***_*n*_ whose entries are nonnegative and sum to one; *θ*_*g*_ is the gene-specific inverse dispersion; and *π*_*ng*_(***z***_*n*_, *b*_*n*_) is the probability of an extra zero. The encoders *q*_*ϕ*_(***z***_*n*_ |***x***_*n*_, *b*_*n*_) and *q*_*ϕ*_(*ℓ*_*n*_ |***x***_*n*_, *b*_*n*_), which infer the two latents from the counts, are trained to maximize the evidence lower bound,

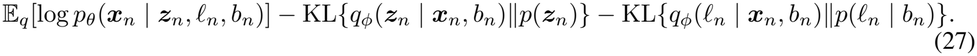

We train this baseline with the exact scPACE training-cell indices and the persisted decoder batch, and we supply neither phenotype nor cell type. We report the posterior-mean cell latent E_*q*_[***z***_*n*_|***x***_*n*_, *b*_*n*_], evaluated by the same encoder for validation and external-query cells. scVI thus provides a vector representation.

#### A.4.8 scANVI

scANVI extends scVI with a cell-type label *c*_*n*_ ∈{1, …, *K}*, where *K* is the number of cell types, and with an auxiliary latent ***u***_*n*_ [Xu et al., 2021]. In their published method, a compact form of the generative model is

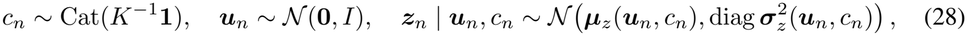

where *c*_*n*_ is drawn from a uniform categorical over the *K* types, meaning each type is equally likely a priori; ***u***_*n*_ is standard normal; and the representation ***z***_*n*_ is Gaussian with mean and variance produced from ***u***_*n*_ and *c*_*n*_. The batch-aware scVI count likelihood then generates the counts from ***z***_*n*_. The inference model has three encoders, *q*_*ϕ*_(***z***_*n*_| ***x***_*n*_), *q*_*ϕ*_(*c*_*n*_ | ***z***_*n*_), and *q*_*ϕ*_(***u***_*n*_ | ***z***_*n*_, | *c*_*n*_). For an unlabeled cell, the training objective averages the labeled-cell objective over the *K* candidate types, weighted by *q*_*ϕ*_(*c*_*n*_ | ***z***_*n*_), and adds the entropy of that type distribution.

We expose only training-cell types during training, and give external-query cells unlabeled category. We report the posterior cell representation together with the full class-probability vector *q*_*ϕ*_(*c*_*n*_ | ***z***_*n*_). scANVI therefore adds cell-type transfer to scVI’s representation, but still models no donor-level phenotype.

#### A.4.9 scVIP

scVIP links single-cell expression and cell-type composition to donor phenotypes through a hierarchical generative model, and it summarizes each donor by pooling that donor’s cells with cell-type-aware attention [Lai et al., 2026]. In their published method, for donor *d* and cell *n*, write *c*_*dn*_ for the cell’s observed type, ***u***_*dn*_ for the auxiliary variable, ***z***_*dn*_ for the cell latent, and ***t***_*d*_ for the donor latent. The generative structure includes

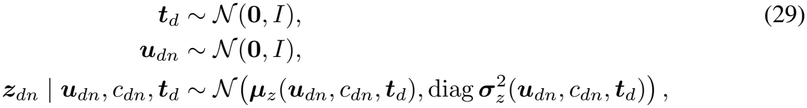

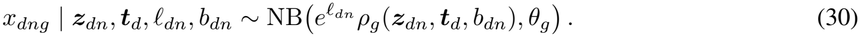

Here the donor latent ***t***_*d*_ and the standard-normal auxiliary variable ***u***_*dn*_ together shape each cell latent ***z***_*dn*_, and the counts *x*_*dng*_ follow a negative-binomial likelihood with library size *ℓ*_*dn*_, batch *b*_*dn*_, and gene-specific inverse dispersion *θ*_*g*_. scVIP ties composition and phenotype to the donor state through a Dirichlet-multinomial model ***N***_*d*_ ∼ DirMult(*n*_*d*_,*φ*_*d*_) with logφ_*d*_ = ***α*** + *B****t***_*d*_, where ***N***_*d*_ is the donor cell type composition;φ_*d*_ contains positive Dirichlet-multinomial concentration parameters, and the normalized concentrations determine the expected cell-type proportions; *B* maps the donor state to them; and ***α*** controls overdispersion. Its cell-level component is initialized from a scVI-family variational autoencoder. To summarize a donor, scVIP pools the cells of each type *k* by gated attention, which learns a weight for each cell and takes a weighted average,

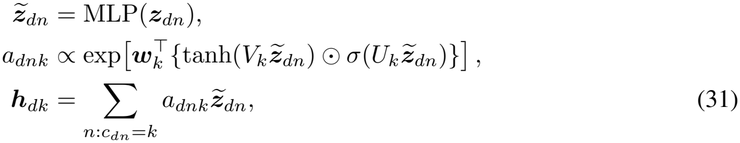

where 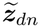 is a transformed cell latent from a small neural network (MLP); ⊙ is the elementwise product; *V*_*k*_, *U*_*k*_, and ***w***_*k*_ are attention parameters; *a* is the attention weight of cell *n* for type *k*; and ***h***_*dk*_ is the resulting type-*k* summary. Concatenating the type summaries {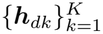 with the donor’s type proportions parameterizes a Gaussian donor posterior *q*_*ϕ*_(***t***_*d*_), and phenotype-specific regression or classification heads model *p*(*y*_*d*_ |***t***_*d*_). The attention weights show which cells and types inform the donor representation.

### A.5 reported scores and confidence threshold

scPACE reports point estimates. It uses the posterior mean of each latent variable as a point value and passes it to the next stage, rather than averaging each reported quantity over the full posterior,

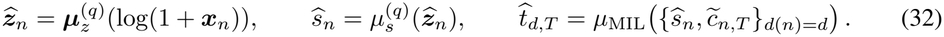

Here 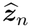 is the cell representation, read as the encoder mean 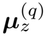 of the log-transformed counts log(1 + ***x***_*n*_ ). The cell score *Ŝ*_*n*_ is read from that representation by the cell-state encoder mean 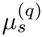. The donor state 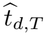 pools the scores *Ŝ*_*n*_ and the transferred types 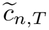 of donor *d*’s cells through the donor-level MIL pooling *µ*_MIL_, where 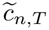 . is the type assigned to a cell only when the classifier is confident at a threshold *T*. The representation 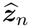 and the cell score *Ŝ*_*n*_ do not depend on *T* . The donor state does, because the pooling is done separately within each type and uses only the cells whose type is confidently assigned at *T* . For a binary donor outcome, the predicted probability is sigmoid 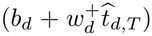, where *b*_*d*_ and 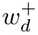 are the bias and positive slope of the donor head. The confidence score 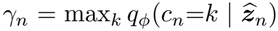 is the largest class probability the classifier assigns to cell *n*. It ranks cells by how certain the classifier is.

### A.6 Simulation data generation and evaluation protocol

The simulation is built from a factorial design over two expression regimes and from a hierarchy of ground truths for donors and cells. The design crosses Regime A and Regime B with *D* ∈ {5, 30, 100} donors and *N*_*d*_ ∈ {100, 1000 }cells per donor at data seed 0, producing 12 separate cohorts and 297,000 cells in total. Every cohort has *G* = 20,000 genes, *K* = 6 abstract cell types, and *R* = 12 latent factors. For donor *d* and cell *n* nested within that donor, the generator draws

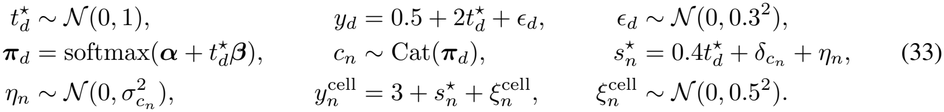

Here *α*_*k*_ ∼ *N*(0, 0.5^2^) and *β*_*k*_ ∼ *N*(0, 0.8^2^). The six type offsets 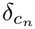 are a seed-fixed permutation of {−2.5, −1.5, −0.5, 0.5, 1.5, 2.5}, and 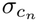 is linearly spaced from 0.20 to 1.10. Structural 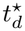 and 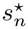 are retained only for evaluation. Fitting receives *y*_*d*_ and the noisy weak-cell target 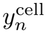.

#### Smooth progression program (Regime A)

For cell *n*, 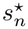 is its scalar ground-truth progression state. The generator transforms this scalar into two fixed features,

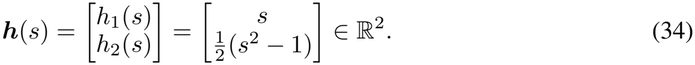

Here, *h*_1_(*s*) = *s* represents a linear progression effect, whereas 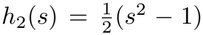 represents a quadratic progression effect. The subtraction of 1 centers the quadratic feature when *s* has unit second moment.

For each gene *g* ∈ {1, …, *G*}, an independent indicator

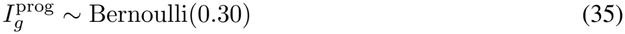

determines whether the gene belongs to the progression-gene pool. Thus, 30% of genes are expected to have a direct progression effect. For every progression gene, the generator draws a shared coefficient vector

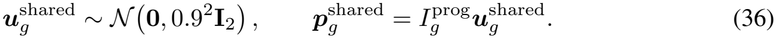

The two entries of 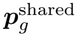 are the shared linear and quadratic coefficients of gene *g*.

Each cell type also has a private progression program. For cell type *k*∈{ 0, …, *K*− 1 }and gene *g*, independently draw

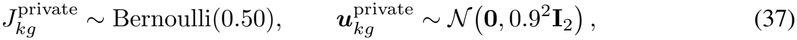

and define

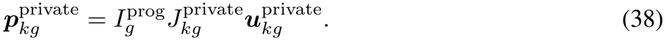

Therefore, the private program of each cell type is restricted to the progression-gene pool and is nonzero for an expected 50% of that pool.

The effective coefficient vector for gene *g* in cell type *k* is

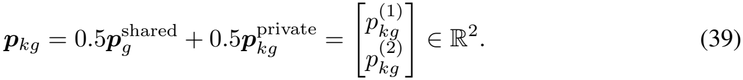

Thus, 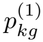 is the linear coefficient and 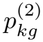 is the quadratic coefficient. Equivalently,

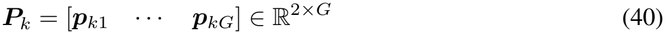

is the complete progression-coefficient matrix for cell type *k*, and ***p***_*kg*_ is its *g*th column. If cell *n* has cell type *c*_*n*_, its smooth progression effect on gene *g* under Regime A is

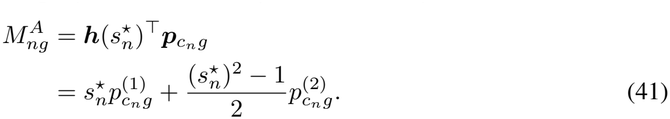

#### Rough progression program (Regime B)

Regime B adds a cell-specific rough effect to an expected 10% of all genes. Because rough genes must belong to the 30% progression-gene pool, the conditional rough-gene indicator is

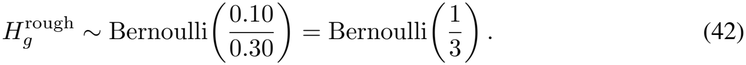

The rough coefficient for gene *g* is

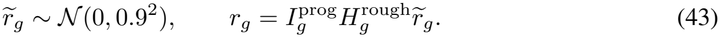

Consequently, *r*_*g*_ = 0 for genes that are not selected as rough genes. Each cell also receives an independent jitter value

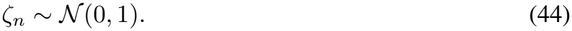

The rough progression effect for cell *n* and gene *g* is therefore

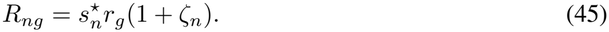

Regime B combines the smooth and rough effects as

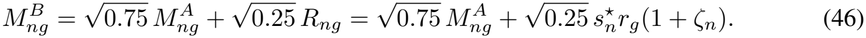

The value 0.25 is a nominal squared-amplitude weight. It does not guarantee that the rough term explains exactly 25% of the realized expression variance. Relative to Regime A, Regime B weakens the smooth program and adds cell-specific variation that depends on 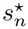 .

#### Gene-expression and count generation

Let *d*(*n*) denote the donor containing cell *n*. For either regime, define 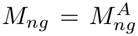 in Regime A and 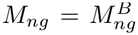 in Regime B. The unnormalized log-expression score of gene *g* in cell *n* is

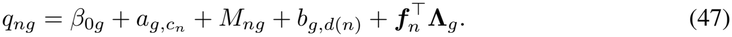

Each term has the following explicit definition.

The gene-specific baseline is

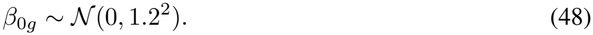

Cell type 0 is the reference type, so

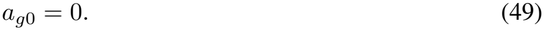

For every nonreference type *k* ∈ {1, …, *K* − 1},

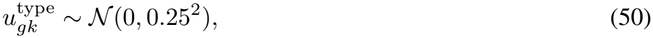

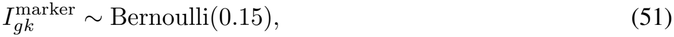

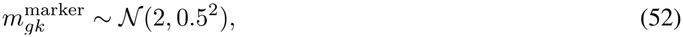

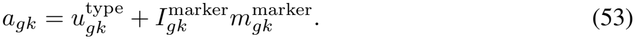

Thus, all nonreference gene-type pairs have a small baseline type effect, and an expected 15% receive an additional marker-gene boost.

For each gene, the donor-effect standard deviation and donor-specific intercepts are

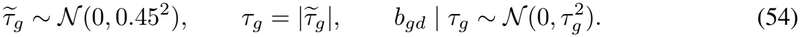

The generator uses *R* = 12 nuisance factors. For cell type *k*, cell *n*, and gene *g*,

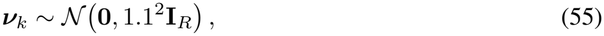

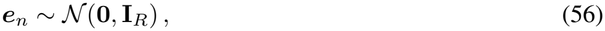

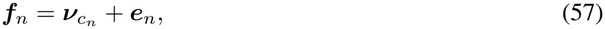

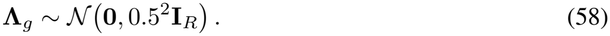

Here, 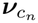 gives cells of the same type a common location in factor space, ***e***_*n*_ adds cell-specific variation, and 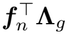 creates correlated expression across genes.

The vector ***q***_*n*_ = (*q*_*n*1_, …, *q*_*nG*_)^*⊤*^ is converted into relative gene-expression probabilities using a softmax over genes:

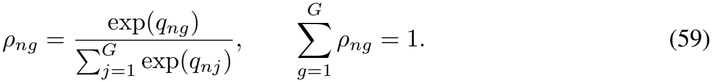

The expected count of gene *g* is

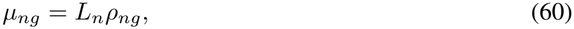

where

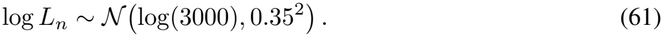

Therefore, *Σ*_*g*_ *µ*_*ng*_ = *L*_*n*_, and *L*_*n*_ is the expected total library size of cell *n*.

To define gene-specific inverse dispersion, first standardize the baseline intercept:

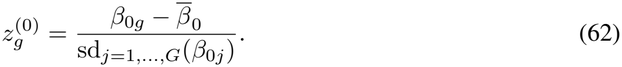

Then draw

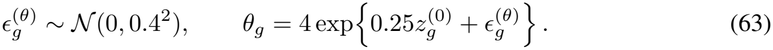

Thus, the inverse dispersion has base scale 4, depends on the gene’s baseline expression, and contains gene-specific log-normal heterogeneity.

Finally, counts are generated through a Gamma–Poisson mixture:

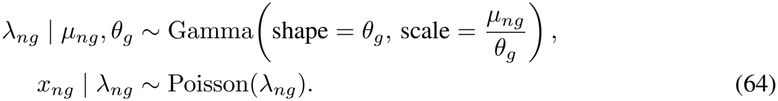

After integrating out *λ*_*ng*_, this is a negative-binomial distribution with

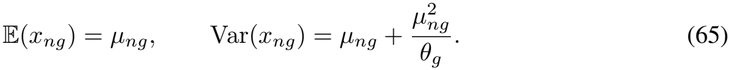

The simulation contains no biological batch effect.

#### Matching Regimes A and B

For a fixed donor count, cells per donor, and data seed, Regimes A and B use the same deterministic random-number streams for all shared quantities. Consequently, the paired regimes have identical donor states 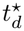, cell states 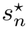, donor and cell phenotypes, cell types, cell-type composition, libraries, nuisance factors, gene baselines, dispersions, donor random effects, and donor partitions. They also share the same smooth coefficient matrices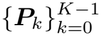.

Regime B alone draws the rough-gene coefficients and cell-specific jitter from separate random-number streams. The regimes therefore differ only in the effective progression term *M*_*ng*_, the resulting expression probabilities and means, and the sampled counts. The 12 factorial settings are generated as separate cohorts; they are not nested subsamples of larger cohorts.

#### Donor partitions

For a cohort with *D* donors, the number of external-query donors is

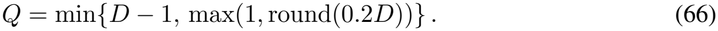

The *Q* donors are sampled without replacement using a deterministic random-number stream derived from data seed 0. This gives *Q* = 1, 6, and 20 for *D* = 5, 30, and 100, respectively.

Among the remaining *D* − *Q* donors, the number of fitting donors is

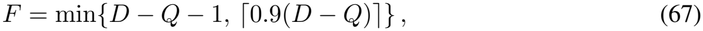

and the number of internal-validation donors is

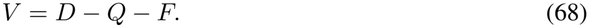

The remaining donors are shuffled using split seed 0; the first *V* donors are assigned to internal validation and the other *F* donors are assigned to model fitting. The resulting fitting/internal-validation/external-query donor counts are 3*/*1*/*1, 22*/*2*/*6, and 72*/*8*/*20 for *D* = 5, 30, and 100.

All methods use these persisted donor roles and the same eligible fitting cells. External-query pheno-types and the structural truths 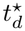 and 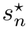 are unavailable during model fitting and score orientation.

We now give the training profile and the evaluation protocol. The matched scPACE profile uses latent width 10, encoder/decoder hidden width 128, one hidden layer, MIL hidden width 32, dropout 0.1, gene chunks of 2,048, and confidence threshold *T* = 0.6. Training runs for at most 400 epochs with shuffled internal splitting, three Monte Carlo validation samples, early stopping with patience 45 and minimum improvement 0, training seed 0, and resumable checkpoints. The loss weights are 1 for reconstruction and each KL term, 400 for donor supervision, 100 for weak-cell supervision, 10 for composition, 50 for type classification, and 0 for the Mundlak term. Optimization uses learning rate 10^*−*3^, gradient clipping at 5, and bfloat16 where supported. Dataset-derived dimensions determine the number of types, decoder batches, and effective genes.

The scalar benchmark compares scPACE with Sceptic, psupertime, PhenoPath, Slingshot, CNA, and MELD. For cell-state recovery, an affine calibration of 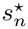 on *ŝ*_*n*_ is fitted using training cells only. External-query performance is summarized by Pearson correlation and

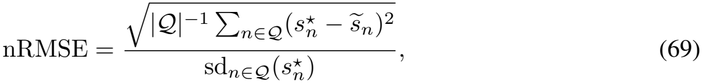

where 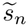 is the training-calibrated score and Q contains external-query cells. For scPACE, the native MIL posterior mean 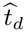 is compared with 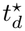 by Pearson correlation and training-calibrated RMSE. The six scalar baselines do not output a native donor state and their *t*-state entries are explicitly not applicable. scVIP held-out prediction uses training-only cell-type imputation before native per-type MIL inference and is compared with 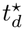 by Pearson correlation. Correlations require at least two external-query donors, and hence are recorded as not estimable for the five-donor cohorts.

Representation comparisons use the scPACE cell embedding together with scVI, scANVI, and scVIP embeddings. All representation probes are fitted on training donors, and are evaluated on externalquery cells or donor aggregates.

### A.7 Real-data cohort construction and evaluation

We first describe the cohorts and how they are harmonized. The integrated study combines 568,654 class-annotated scRNA-seq cells from 53 animals in the Gao et al. developing-mouse visual-cortex atlas with 296,358 project-processed snRNA-seq profiles from 35 GSE190940 technical-library/sample units [Gao et al., 2025, Cheng et al., 2022]. The resulting matrix contains 865,012 profiles over 30,566 shared genes. Gao et al. spans embryonic through adult development and supplies a 15-class vocabulary; GSE190940 samples postnatal development under normally reared, dark-reared, and dark-light-re-exposed conditions. Dataset identity is the two-level decoder batch, while the original library identifier is retained as the sample unit. Gao cell class labels are observed, and every GSE190940 cell class label is treated as missing during training and evaluation.

We construct two phenotype targets, a normalized age at donor level and a maturity score at cell level. At the donor level, age labels were converted to days relative to birth and linearly mapped to [−1, 1] using the range in the 67-unit pre-query reference pool (from −7.5 to 68 days). At the cell level, using AUCell [Aibar et al., 2017, Van de Sande et al., 2020], we constructed a predefined AUCell-like mature-minus-progenitor score on the raw-count matrix. Let *G* = 30,566 be the number of shared genes and *K* = max {1, round(0.05*G*) }= 1,528 be the rank cutoff used by the AUCell-like statistic. Within each cell, genes were ordered by decreasing count after a seeded permutation that resolves ties deterministically (seed 0). If *r*_*ng*_ ∈ {0, …, *G* − 1} is the resulting rank of gene *g* in cell *n*, with 0 denoting the highest expression, the score for marker set *S* is

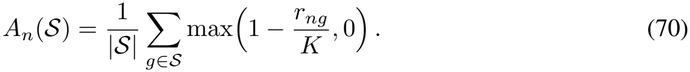

The retained mature set *ℳ* comprised *Rbfox3, Satbf2, Cux2, Mef2c, Tbr1, Snap25, Syt1, Grin1, Camk2a*, and *Mapt*; the requested marker *Bcl11b* was absent from the shared feature space. The progenitor set *P* comprised *Sox2, Pax6, Hes1, Hes5, Nes, Vim, Mki67, Top2a, Ccnd2, Eomes, Neurog2*, and *Btg2*. Let *ℛ* contain all cells from the 67 pre-query reference units and let *z*_*ℛ*_ (*v*_*n*_) = {*v*_*n*_ − *µ*_*ℛ*_ (*v*)}*/σ*_*ℛ*_ (*v*) denote standardization using the population mean and standard deviation in *ℛ*. The final maturity target is

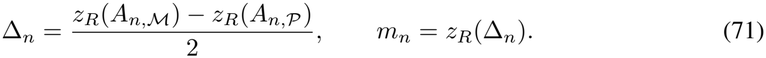

Marker coverage was determined from the final shared feature space, whereas all age and maturity centering and scaling constants were estimated from the pre-query reference pool. No external-query cell or unit was used to estimate these constants. Only training cells contribute target gradients.

For evaluation, all non-normally-reared GSE190940 samples and 10% of normally reared samples in each dataset were assigned to the external query. This yields 21 query samples and 183,656 cells: 6 Gao et al. samples (52,814 cells) and 15 GSE190940 samples (130,842 cells). The remaining 67 samples were split into 61 training samples (621,752 cells) and 6 internal-validation samples (59,604 cells). Every benchmark uses the same 61 training units, and external-query phenotypes do not enter fitting, score orientation, or calibration.

For the model configuration, the real-data scPACE profile uses a 10-dimensional cell latent, hidden width 128, one hidden layer, MIL width 32, dropout 0.1, gene chunks of 2,048, at most 400 epochs, training seed 0, and confidence threshold *T* = 0.6. Reconstruction and KL weights are 1; donor, cell, composition, and class weights are 400, 100, 10, and 50.

We report the recovery metrics and secondary results next. Cell-state recovery is Pearson correlation between a method’s reported scalar output and the maturity AUCell score on all 183,656 external-query cells. scPACE reaches *r* = 0.919 overall, with *r* = 0.962 on Gao cells and *r* = 0.844 on GSE cells. Sample-level evaluation compares scPACE 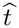 and the scVIP phenotype prediction with observed normalized age on the same 21 query units. Their Pearson correlations are 0.942 for scPACE and 0.542 for scVIP, respectively.

Raw-argmax cell-class accuracy is evaluated only on the 52,814 typed Gao query cells. scPACE and scANVI attain accuracies 0.9772 and 0.9765.

Donor mixing is evaluated using the within-class 15-nearest-neighbor different-donor fraction, divided by its random-label expectation. Computed on a 9,998 cell Gao et al. subset. We have 0.1377, 0.1317, 0.1395, and 0.1365 for scPACE, scVIP, scANVI, and scVI.

Finally, we describe the visualization and the condition diagnostic. Figure 2e,f uses one UMAP of the scPACE latent embedding and renders all 865,012 cells without display downsampling. In cell class UMAP, colors for Gao et al. data are observed cell classes, whereas GSE190940 colors are raw scPACE argmax predictions.

